# Modulation of the Drosophila Transcriptome by Developmental Exposure to Alcohol

**DOI:** 10.1101/2021.12.09.472007

**Authors:** Tatiana V. Morozova, Vijay Shankar, Rebecca A. MacPherson, Trudy F. C. Mackay, Robert R. H. Anholt

## Abstract

Prenatal exposure to ethanol can cause fetal alcohol spectrum disorder (FASD), a prevalent, preventable pediatric disorder. Identifying genetic risk alleles for FASD is challenging since time, dose, and frequency of exposure are often unknown, and manifestations of FASD are diverse and evident long after exposure. *Drosophila melanogaster* is an excellent model to study the genetic basis of the effects of developmental alcohol exposure since many individuals of the same genotype can be reared under controlled environmental conditions. We used 96 sequenced, wild-derived inbred lines from the *Drosophila melanogaster* Genetic Reference Panel (DGRP) to profile genome-wide transcript abundances in young adult flies that developed on ethanol-supplemented medium or standard culture medium. We found substantial genetic variation in gene expression in response to ethanol with extensive sexual dimorphism. We constructed sex-specific genetic networks associated with alcohol-dependent modulation of gene expression that include protein-coding genes, Novel Transcribed Regions (NTRs, postulated to encode long non-coding RNAs) and female-specific coordinated regulation of snoRNAs that regulate pseudouridylation of ribosomal RNA. We reared DGRP lines which showed extreme upregulation or downregulation of snoRNA expression during developmental alcohol exposure on standard or ethanol supplemented medium and demonstrated that developmental exposure to ethanol has genotype-specific effects on adult locomotor activity and sleep. There is significant and sex-specific natural genetic variation in the transcriptional response to developmental exposure to ethanol in Drosophila that comprises networks of genes affecting nervous system development and ethanol metabolism as well as networks of regulatory non-coding RNAs.

**Summary statement:** We developed a Drosophila model for Fetal Alcohol Spectrum Disorder and found that developmental exposure to alcohol modulates expression of networks of genes affecting nervous system development and ethanol metabolism as well as networks of regulatory noncoding RNAs.

## Introduction

Prenatal exposure to ethanol can trigger a wide range of adverse physiological, behavioral, and cognitive outcomes, referred to as fetal alcohol spectrum disorder (FASD) [1–4]. Fetal alcohol syndrome (FAS) is the most severe FASD, presenting with craniofacial dysmorphologies, neurocognitive deficiencies, and behavioral disorders, including hyperactivity, attention deficit disorder and motor coordination anomalies [1,5–7]. FAS/FASD is the most common preventable pediatric disorder, often diagnostically confounded with autism spectrum disorder [8]. The Centers for Disease Control and Prevention found that, despite warning labels on alcoholic beverages that indicate possible adverse effects on prenatal development, 1 in 10 pregnant women report alcohol use and more than 3 million women in the USA are at risk of exposing their developing fetus to alcohol [9]. Although defects from prenatal alcohol exposure can be replicated in mouse models [10], identifying genetic factors that contribute to susceptibility to FASD is virtually impossible in human populations since time, dose, and frequency of exposure are often unknown, and manifestations of FASD are diverse and become evident long after exposure.

*Drosophila melanogaster* is an excellent model to study the genetic and genomic bases of the effects of developmental alcohol exposure. The Drosophila model allows strict control over the genetic background, and we can rear large numbers of individuals of the same genotype under controlled environmental conditions without regulatory restrictions and at low cost. In addition, following acute exposure to alcohol, flies undergo physiological and behavioral changes that resemble human alcohol intoxication, including loss of postural control, sedation, and development of tolerance [11–14].

Previous studies on the effects of developmental alcohol exposure in Drosophila showed reduced viability and delayed development time [15,16], reduced adult body size [15], and disruption of neural development [17]. Developmental exposure to alcohol was associated with reduction in the expression of a subset of insulin-like peptides and the insulin receptor [15], dysregulation of lipid metabolism and concomitant increased oxidative stress [18] and reduced larval food intake due to altered neuropeptide F signaling [19].

In previous studies, we have taken advantage of natural variation in the *Drosophila melanogaster* Genetic Reference Panel (DGRP), a well characterized population of 205 inbred wild-derived lines with complete genome sequences [20,21], to study the genetic underpinnings of developmental alcohol exposure [16], voluntary ethanol consumption [22], acute ethanol intoxication, and induction of tolerance [23,24], a prelude to the development of alcohol dependence in people. Linkage disequilibrium decays rapidly within a few hundred base pairs in this population [21], enabling identification of candidate causal SNPs associated with phenotypic variation in alcohol-related traits. A major unresolved question relates to the mechanism(s) by which alcohol exposure during development affects adult phenotypes. Here, we performed RNA sequencing to assess the effects of developmental alcohol exposure on genome wide genetic variation in gene expression of young adults.

## Results

### Transcriptional profiles of flies reared on ethanol supplemented medium

We performed transcriptional profiling of flies reared on regular medium or medium supplemented with 10% (v/v) ethanol from 96 DGRP lines that span the phenotypic spectrum of alcohol sensitivity following developmental exposure to alcohol (Additional File 1) [16]. For each line, we obtained duplicate samples of 30 males and 25 females, aged 3-5 days, all collected at the same time of day, and performed RNA sequencing using 125 bp single end reads. A total of 33,580 transcripts were expressed in adult flies, of which 16,165 (48.1%) are annotated and 17,415 (51.9%) are novel transcribed regions (NTRs) [25,26] (Additional file 2). We performed three-way factorial mixed model ANOVAs with the main effects of DGRP line, sex, and treatment for all expressed transcripts, and used a False Discovery Rate (FDR) threshold < 0.05 for statistical significance of each term in the ANOVA model (Additional File 2). As in previous analyses of genome-wide gene expression using whole flies [25–27], we find that expression of nearly all of the expressed transcripts is sexually dimorphic (28,343, 84.4%). The expression of 16,278 transcripts (48.5%) is modulated by alcohol, and for 10,002 transcripts the transcriptional response to ethanol differs between males and females (*i.e.,* there is a treatment by sex interaction). There is significant genetic variation in expression for 10,620 transcripts; as well as context-specific genetic variation, with 11,338 transcripts exhibiting genetic variation in sexual dimorphism (sex by line interaction), 1,222 showing genetic variation in response to developmental exposure to ethanol (treatment by line interaction) and 77 with genetic variation in sexual dimorphism in response to ethanol (sex by treatment by line interaction).

Because we found extensive interactions with sex and treatment and sex and DGRP line, we performed two-way ANOVAs partitioning gene expression variation by line, treatment, and the line by treatment interaction (*L×T*), separately for males and females (Additional File 3). The main effect of treatment was significant (FDR < 0.05) for 14,158 (13,827) transcripts in females (males), and the main effect of line was significant for 13,521 (20,996) transcripts in females (males). We are most interested in transcripts with a significant genotype by alcohol treatment interaction (*L×T*), since these transcripts show genetic variation in their response to developmental ethanol exposure. We found 939 significant *L×T* interactions in females and 823 in males when we compared expression profiles of flies grown on ethanol supplemented medium with transcript abundance levels obtained under standard growth conditions (Additional File 4). Of these 1,351 transcripts that have genetically variable responses to ethanol exposure during development, 253 are in common between males and females, 499 are female-specific and 346 are male-specific. The transcripts with significant *L×T* terms in females were enriched for biological process gene ontology terms involved in metabolism, biosynthesis, transcription, immune/defense response and chromatin organization; and the transcripts with significant *L×T* terms in males were enriched for biological process gene ontology terms involved in immune/defense response, metabolism and development [28] (Additional File 4). A total of 65.6% of these genes have human orthologs (Additional File 4).

### Co-regulated modules of transcripts that are differentially regulated after developmental alcohol exposure

We analyzed the sex-specific correlation structure of the developmental alcohol-sensitive transcriptome and identified eight highly interconnected modules in females (Fig. 1). These modules contained transcripts associated with xenobiotic detoxification and metabolism (Figs. 1 A, B), development and cell adhesion (Figs. 1B, C), cuticle formation (Fig. 1E) and neural signaling (Fig. 1G). The latter includes *sNPF,* associated with neuropeptide F signaling previously implicated with reduced larval food intake when larvae were grown on alcohol-supplemented medium [19]. One module consists almost exclusively of NTRs (designated XLOCs; Fig. 2D) [25,26]. Of special interest is *Ilp3,* which encodes an insulin-like peptide implicated in developmental exposure to alcohol [15] and is correlated with a network of NTRs (Fig. 1F). The most highly correlated group of transcripts are the snoRNAs (Fig. 1H).

**Figure 1.**
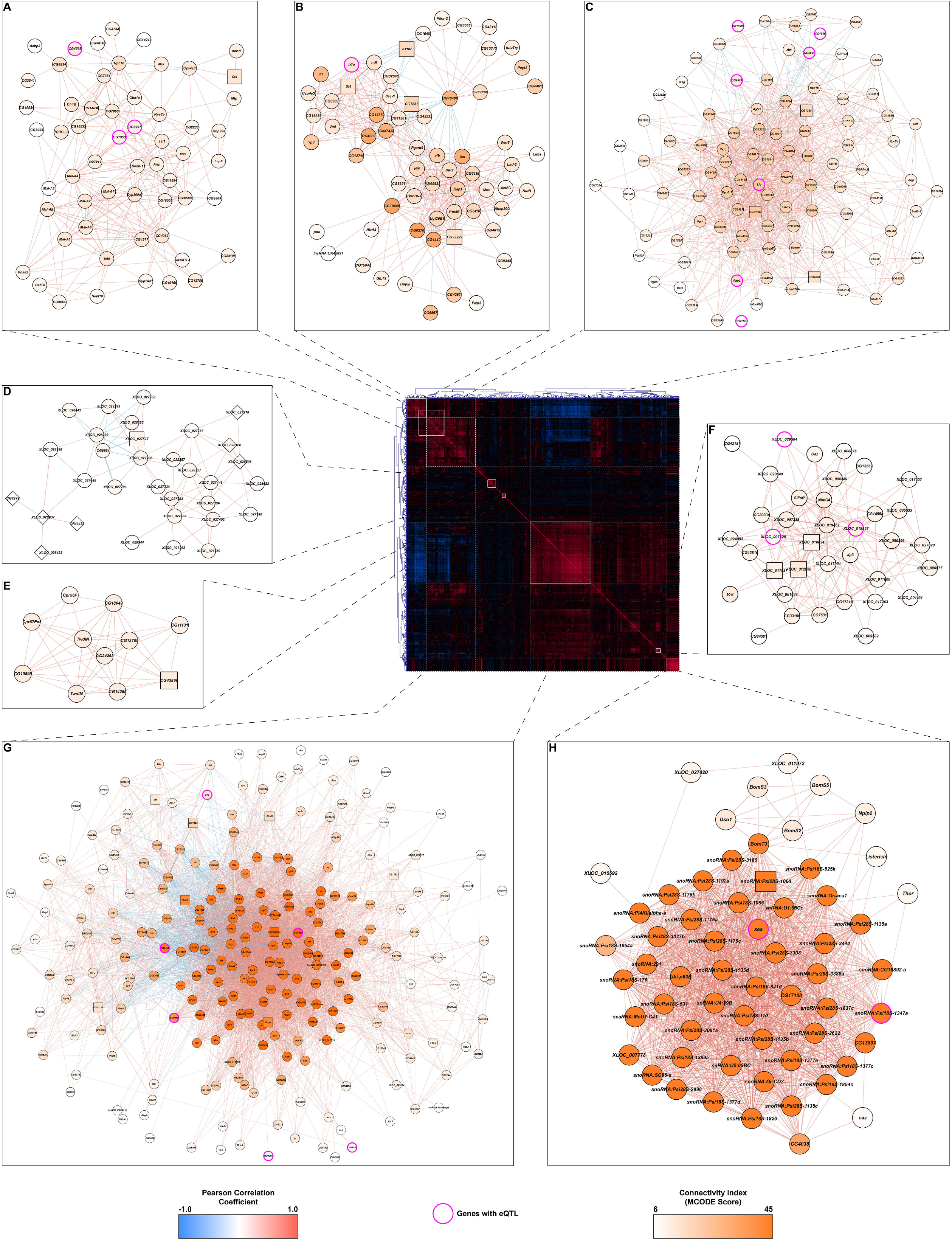
Correlations of differences in gene expression between developmental ethanol treatment and control in females. The center panel heat map corresponds to the unfiltered, bi-clustered correlation matrix calculated for differences in expression of genes with a statistically significant line-by-ethanol treatment interaction term in a linear mixed effects model. The strength of the correlation is depicted as gradients and the directionality as color (positive correlations in red and negative correlations in blue). Networks derived from clusters with strong intra-connectivity are depicted around the center panel (panels **A-H**). The MCODE connectivity score for each node is represented as a color gradient. The edge colors follow the same scheme as the center panel (strength as gradient and directionality as color). Genes with statistically significant eQTLs are highlighted with pink borders.

Although there was overlap between alcohol-modulated transcripts in males and females, the correlation structure for differentially expressed genes in males shows an entirely different modular organization (Fig. 2). The nine male modules are generally smaller and more difficult to interpret than the female modules, as they are dominated by NTRs and genes of unknown function. Whereas the functions of this new class of non-coding transcripts remain to be established, they feature prominently in networks of alcohol-modulated transcripts (Figs. 1 and 2), suggesting a regulatory role for non-coding elements in the genome in modulating the transcriptional response to developmental alcohol exposure.

**Figure 2.**
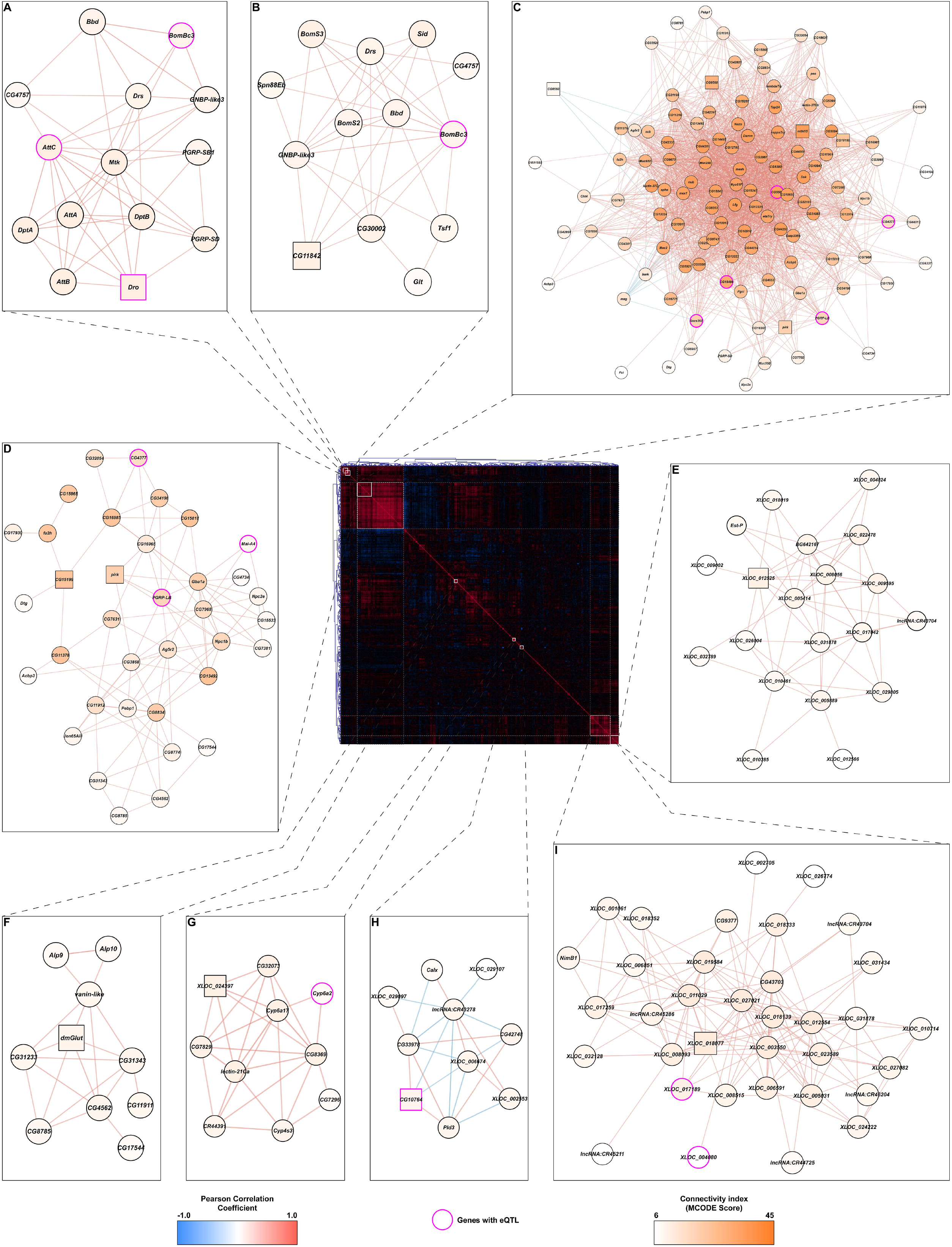
Correlations of differences in gene expression between developmental ethanol treatment and control in males. The center panel heat map corresponds to the unfiltered, bi-clustered correlation matrix calculated for differences in expression of genes with a statistically significant line-by-ethanol treatment interaction term in a linear mixed effects model. The strength of the correlation is depicted as gradients and the directionality as color (positive correlations in red and negative correlations in blue). Networks derived from clusters with strong intra-connectivity are depicted around the center panel (panels **A-I**). The MCODE connectivity score for each node is represented as a color gradient. The edge colors follow the same scheme as the center panel (strength as gradient and directionality as color). Genes with statistically significant eQTLs are highlighted with pink borders.

To identify genetic associations within modules of differentially expressed genes, we mapped expression quantitative trait loci associated with the difference in expression between standard and ethanol-supplemented medium, or response e-QTLs [25,26]. We identified 53 eQTLs, including 19 NTRs, in females, and 45 eQTLs, including 11 NTRs and four long noncoding RNAs (lncRNAs), in males (Additional File 5). All of these eQTLs, with the exception of the eQTL associated with the difference in *NetA* expression between standard and ethanol-supplemented medium in females, were in *trans* (greater than 1 kb from the start and end of the gene body) to the genes associated with an *LxT* interaction). Therefore, we determined to what genes these eQTLs were in *cis* to (defined by within 1 kb of a gene body) (Additional File 5). We then input the genes with *L*x*T* interactions and the genes to which the eQTLs are *cis* into known genetic and protein-protein or RNA-protein physical interaction networks, to construct sex-specific networks associated with genetic variation in response to developmental exposure to ethanol (Fig. 3). The resulting networks are composed of integrative modules that highlight considerable overlap between cellular processes in males and females despite significant sexual dimorphism in ethanol-induced differential gene expression. Genes associated with neuronal development and differentiation, response to ethanol, and reproduction are evident in networks of both sexes. The female network also incorporates modules associated with glutathione metabolism and phototransduction (Fig. 3A), whereas the male network contains modules associated with immune response, starvation and stress response, and septate junction assembly (Fig. 3B).

**Figure 3.**
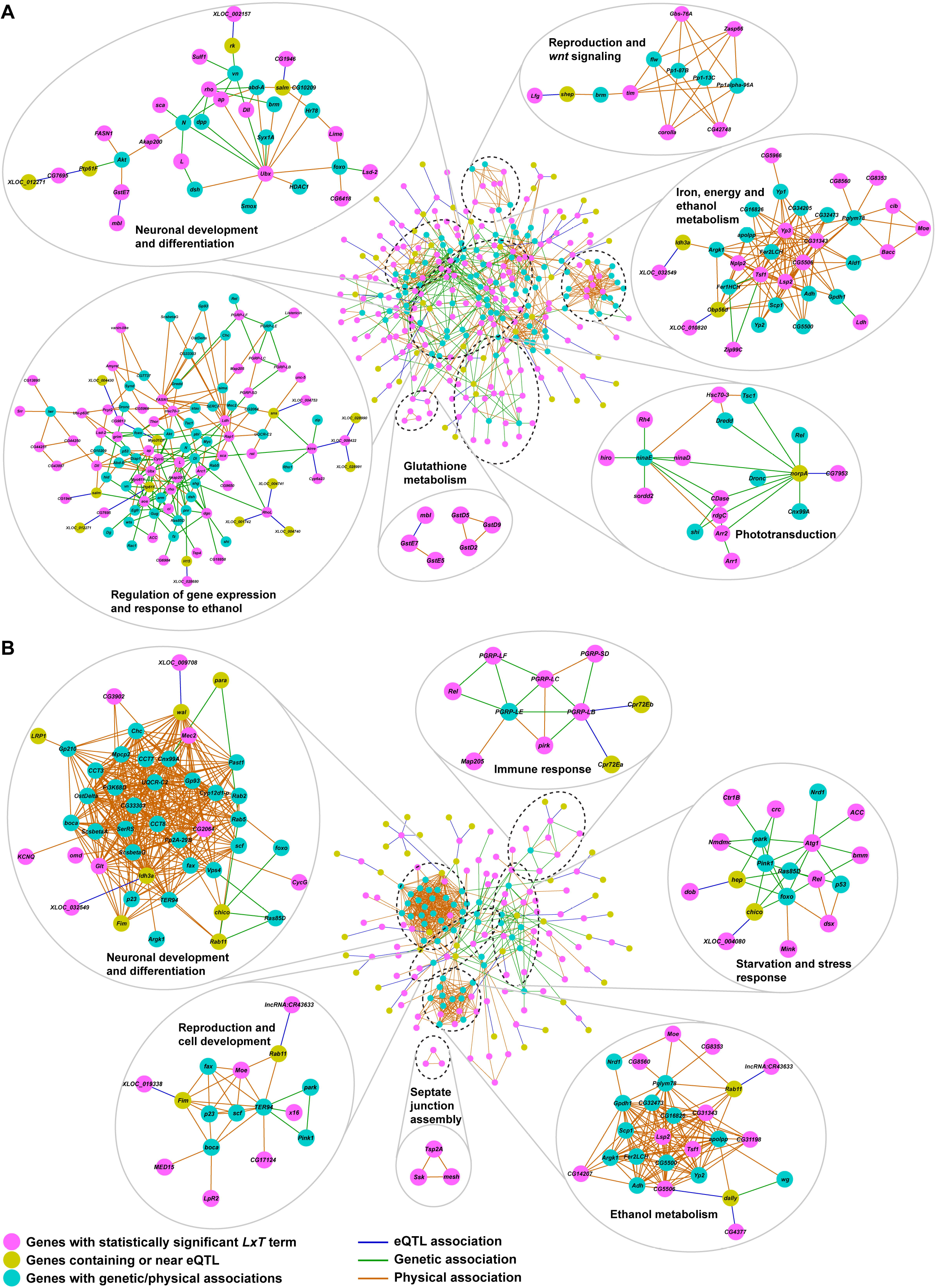
Female (A) and male (B) interaction networks built from eQTLs and known genetic and physical associations. The central networks in each panel represent the sex-separated, filtered interaction networks generated by incorporating eQTL associations calculated from expression differences, between ethanol and control conditions, of genes with a statistically significant line-by-treatment (*LxT)* term in the ANOVA model, to known genetic and protein-protein or RNA-protein physical interactions from the FlyBase interaction database. Pink nodes represent the genes from the *LxT* set. Yellow nodes represent genes either containing or within 1,000 bp of the eQTL variant. Cyan nodes represent genes with known genetic or physical interactions to the rest of the network. Blue edges represent the eQTL associations from this study. Green and orange edges represent known genetic and physical associations from the Flybase interaction database. Individual inlets of genes around the central network are MCODE-generated modules of genes. Annotations of the inlets are based on statistically enriched pathways for genes within these modules. Terms with Benjamini-Hochberg FDR adjusted *P*-value < 0.05 in the statistical overrepresentation test were considered statistically significant.

### Coordinated sex-specific modulation of an ensemble of snoRNAs

Altered co-regulation of 38 snoRNAs was observed only in females exposed to alcohol during development. The direction of changes in snoRNA expression was strongly background-dependent, with some DGRP lines showing coordinated up-regulation, some exhibiting coordinated down-regulation, and others showing no change in snoRNA expression (Fig. 4; Additional File 6). The snoRNAs that exhibit genetic variation in their response to developmental ethanol exposure belong primarily to the H/ACA class, associated with pseudouridylation of rRNAs [29,30], and many are in introns of *RpS4, RpL5, RpS5a, RpS7, RpL11, RpS16, RpL17,* and *RpL22* ribosomal protein encoding genes. The number of *snoRNAs* within each gene varies from 1 to 15 (Additional File 6). When multiple *snoRNAs* are present in a gene, only some show altered expression in response to ethanol exposure, although in some cases clusters of genes, likely expressed as polycistronic transcripts, are regulated together. Other snoRNAs with altered expression in response to chronic ethanol exposure are in introns in *dom,* which contributes to histone acetyl transferase activity associated with epigenetic modification of gene expression [31]; *CG13900 (Sf3b3*), inferred to form part of a spliceosome complex [32]; *SC35,* which encodes a splicing factor [33]; *kra,* which is annotated as a translation initiation factor binding protein [34]; *Aladin,* predicted to form part of the nucleopore complex [35]; and *Nop60B,* which encodes pseudouridine synthase [36,37].

**Figure 4.**
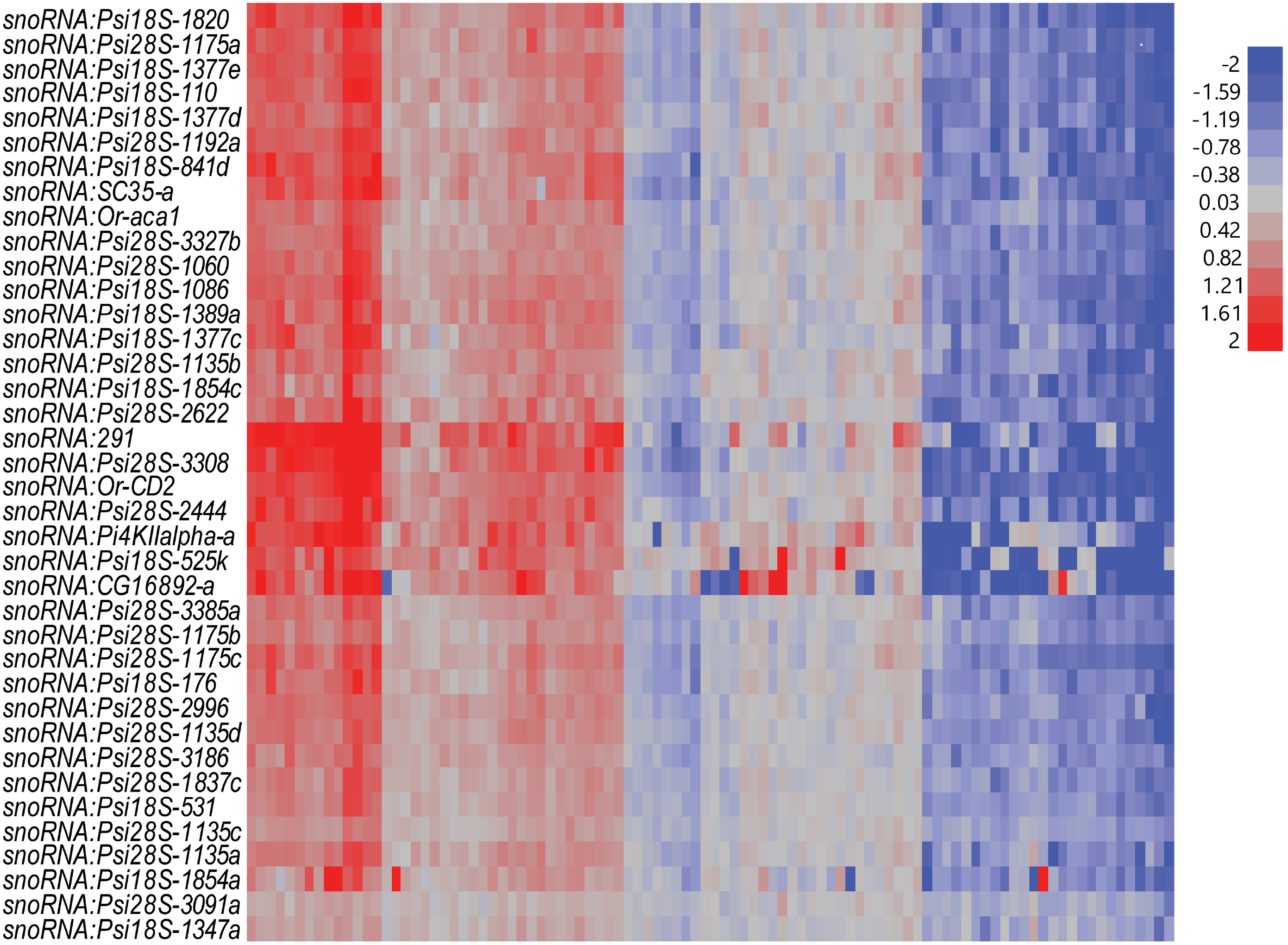
Differentially expressed snoRNAs from female flies grown on ethanol versus regular food. Vertical columns represent individual DGRP lines. The color scale indicates upregulation (red) or down-regulation (blue) as a result of growth on ethanol-supplemented medium.

Examination of variation in transcript abundances of snoRNAs and host genes other than those encoding ribosomal proteins show that the expression of these snoRNAs is regulated independently from the host genes (Fig. S1). Interestingly, transcript abundance levels of all alcohol-sensitive snoRNAs were correlated with variation in expression of *Uhg4,* which is highly expressed in ovaries and encodes a lncRNA (Fig. S1) [38]. A previous genome-wide association study in the DGRP identified *Cyclin E (CycE)* as a highly interconnected hub gene in a genetic network associated with alcohol-induced variation in viability and development time [16]. Variation in *CycE* transcript abundance was not correlated with variation in transcript abundance levels of snoRNAs but was highly correlated with variation in transcript abundance levels of their host genes (Fig. S2). Expression levels of *stet, dom, Aladin, kra* and *Nop60B* were also highly correlated (Fig. S2).

### Effects of developmental alcohol exposure on activity and sleep

The transcriptome is a proximal determinant of organismal phenotypes. FAS/FASD symptoms include hyperactivity and sleep disorders. Since activity and sleep are universal measures of nervous system function which can be modeled in Drosophila, we used activity and sleep parameters as a read-out of the behavioral effects of alcohol exposure during development (Fig. 5). We selected three DGRP lines which exhibited the highest degree of coordinated up-regulation (DGRP_177, DGRP_208, DGRP_367) and down-regulation (DGRP_555, DGRP_705, DGRP_730) of a subset of snoRNAs in response to developmental alcohol exposure and reared them either on regular food or food supplemented with ethanol. We measured their activity and sleep phenotypes using the Drosophila Activity Monitor (DAM) system, in which single flies are introduced into narrow tubes and a movement is recorded any time the fly disrupts an infrared beam. We performed factorial mixed model ANOVA analyses with the main effects of sex, treatment and DGRP line for activity, night and day sleep proportion, and night and day bout count. There was significant genetic variation for all traits among the DGRP lines, and the *L*×*T* interaction was significant for activity, night bout count, and the proportion of day and night sleep (Additional File 7). Thus, exposing Drosophila to alcohol during development affects behavioral phenotypes relevant to patients with FASD, but the effects are genotype specific.

**Figure 5.**
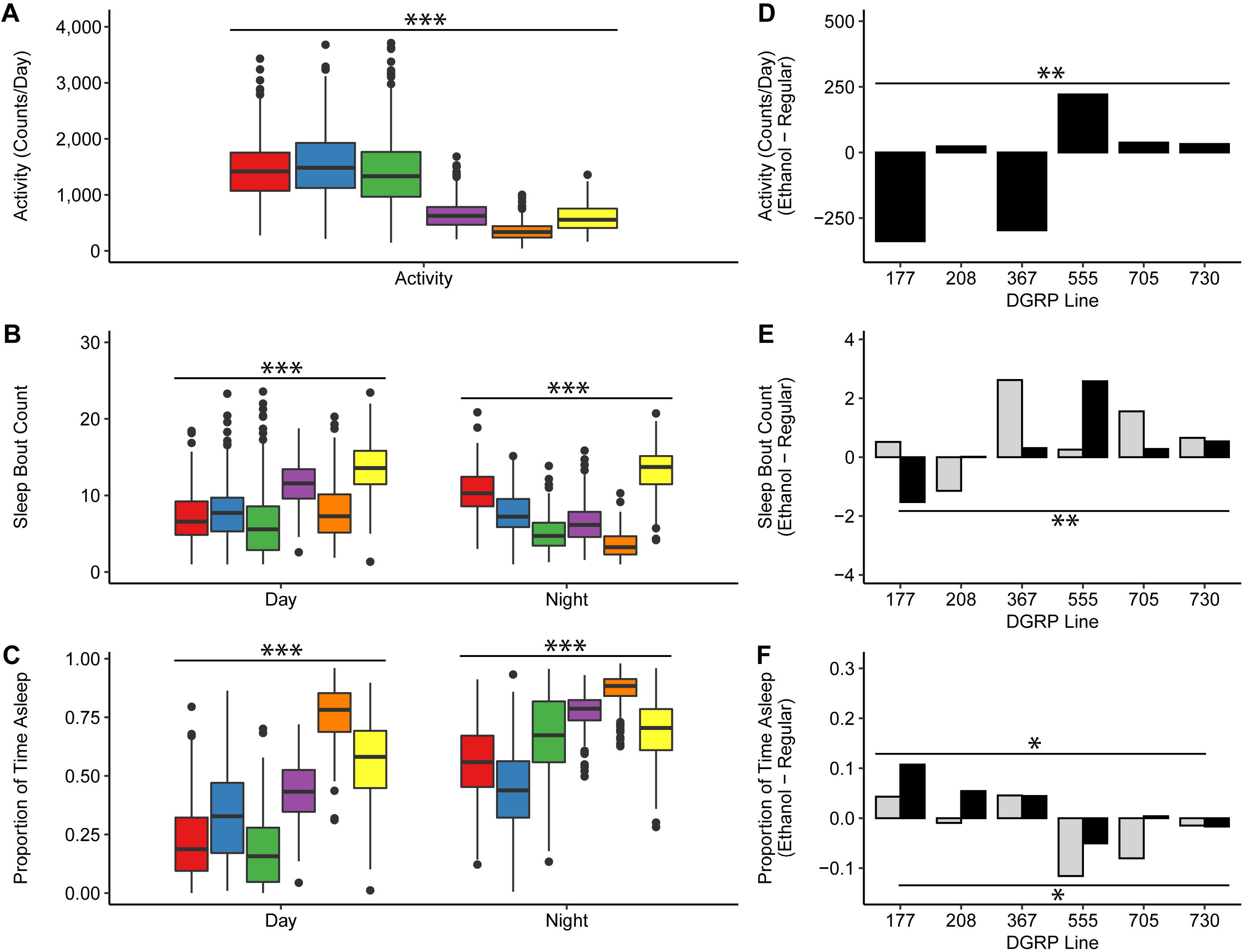
Effects of developmental ethanol exposure on sleep and activity phenotypes. Boxplots averaged across treatment and sex showing the main effect of line on (**A**) locomotor activity, recorded as the average number of counts per day, where counts are the number of times the fly crosses the infrared beam as recorded by the DAM System, (**B**) number of sleep bouts during day and night hours, and (**C**) proportion of time spent asleep during day and night hours. DGRP lines 177, 208, 367, 555, 705, 730 are shown in red, blue, green, purple, orange, and yellow, respectively. Bar graphs of (**D**) locomotor activity (*L*x*T P* = 0.0091), (**E**) number of sleep bouts during day (light grey bars) and night (black bars) hours (Day *L*x*T P* = 0.1538; Night *L*x*T P* = 0.0014), and (**F**) proportion of time asleep during day (light grey bars) and night (black bars) hours (Day *L*x*T P* = 0.0491; Night *L*x*T P* =0.0181), for each DGRP line, averaged across treatment, showing the effect of line by treatment (ethanol-supplemented food minus regular food). * *P* < 0.05, ** *P* < 0.01, *** *P* < 0.001.

## Discussion

Although previous studies have identified a plethora of genetic risk factors that contribute to alcohol related phenotypes in human populations or rodent models [11,39], our understanding of the interaction between environmental alcohol exposure and allelic variants remains incomplete. Evolutionary conservation of fundamental biological processes and similarity of the effects of alcohol exposure between flies and people have established *D. melanogaster* as a useful translational gene discovery system [11,23,40]. Here, we identified transcripts that undergo altered regulation when flies are reared on ethanol. Changes in transcript abundance patterns are sexually dimorphic and reveal regulatory networks in which NTRs feature prominently. It is of special interest that females exposed to developmental ethanol exposure show altered regulation of a large ensemble of H/ACA class snoRNA genes, which may mediate widespread changes in protein synthesis upon chronic alcohol exposure. The complex relationship between expression of host genes and embedded snoRNAs, which we observe in our studies, has also been documented across seven human tissues, including brain [41]. The mechanisms by which alcohol triggers changes in gene expression remain unknown and could include direct effects on the genome, indirect effects on the genome mediated via alcohol-induced metabolic changes, and/or epigenetic modifications of DNA. Pseudouridylation of mRNAs, tRNAs and other small RNAs in response to environmental stress can consolidate or destabilize interactions between RNAs and proteins [42]. The intimate relationship between snoRNAs and ribosomal function may represent a conduit between the genome and the proteome that can adaptively modulate the composition of the proteome in response to ethanol exposure.

The *D. melanogaster* transcriptome is highly intercorrelated [25] and changes in gene expression due to an environmental disturbance result in modulation of transcriptional niches (*i.e.* coregulated ensembles) of focal genes [40,43]. This raises a central “cause *versus* effect” question as it is not evident *a priori* which gene is the focal gene that directly responds to the environmental perturbation. Insights can be derived from eQTL analyses that can delineate *cis*-eQTLs and *trans*-eQTLs [25, 26]. Interestingly, all but one of the eQTLs associated with transcripts with genetic variation in the difference in expression between standard rearing conditions and developmental exposure to ethanol were in *trans* to the focal transcripts. However, we were able to incorporate the genes to which these eQTLs are *cis* into known interaction networks to derive sex-specific networks associated with genetic variation in response to developmental exposure to ethanol in which these genes are candidate regulatory drivers.

The data we present were obtained from flies that were continuously exposed to alcohol from egg to adult. Exposures that are restricted to different developmental stages will provide a finer grained picture of the dynamics of the alcohol-sensitive genome. Similarly, we analyzed transcriptional responses in whole flies. Single cell RNA sequencing experiments with defined tissues, such as the brain, can provide tissue-specific resolution of the transcriptional response to developmental alcohol exposure [44]. Nevertheless, the data we obtained in this study underscore extensive sexual dimorphism and emphasize the importance of non-coding elements in regulating the transcriptional response to alcohol exposure during development. Results from this study illustrate the power of the Drosophila model as a gene discovery system to gain insights into human disorders that can only be addressed through comparative genomics approaches, such as FASD.

## Materials and Methods

### Drosophila lines

We selected 96 lines of the *Drosophila melanogaster* Genetic Reference Panel (DGRP) [20,21] across the range of phenotypic variation of effects of alcohol exposure on viability and developmental time (Additional File 1) [16,24] and reared them on cornmeal-molasses-agar medium supplemented with10% (v/v) ethanol at 25°C, 60-75% relative humidity and a 12-hr light-dark cycle at equal larval densities. We collected two replicates of mated 3-5-day old flies (25 females and 30 males per line) for a total of 384 samples, following procedures described previously for baseline sample collection [26]. We used a randomized experimental design for sample collection that was done strictly between 1-3 pm and froze collected flies over ice supplemented with liquid nitrogen. The flies were sexed and stored in 2.0 ml nuclease-free microcentrifuge tubes (Ambion) at −80°C until processing.

### RNA sequencing

Total RNA was extracted as described previously [26]. Briefly, total RNA was extracted with Trizol using the RNeasy Mini Kit (Qiagen, Inc.) and ribosomal RNA was depleted from 5 ug of total RNA using the Ribo-Zero™ Gold Kit (Illumina, Inc.). Depleted mRNA was fragmented and converted to first strand cDNA. During the synthesis of second strand cDNA, dUTP instead of dTTP was incorporated to label the second strand cDNA. cDNA from each RNA sample was used to produce barcoded cDNA libraries using NEXTflex™ DNA Barcodes (Bioo Scientific, Inc.) with an Illumina TruSeq compatible protocol. Libraries were size-selected for 250 bp (insert size ~130 bp) using Agencourt Ampure XP Beads (Beckman Coulter, Inc.). Second strand DNA was digested with Uracil-DNA Glycosylase before amplification to produce directional cDNA libraries. Libraries were quantified using Qubit dsDNA HS Kits (Life Technologies, Inc.) and Bioanalyzer (Agilent Technologies, Inc.) to calculate molarity. Libraries were then diluted to equal molarity and re-quantified. Samples were processed in batches of 48 and 16 libraries were pooled randomly into 25 pools. Pooled library samples were quantified again to calculate final molarity and then denatured and diluted to 14pM following Illumina guidance. Pooled library samples were clustered on an Illumina cBot; each pool was sequenced on one lane of an Illumina Hiseq2500 using 125 bp single-read v4 chemistry.

### RNA sequence analysis

Sequences were analyzed exactly as described previously [26]. Barcoded sequence reads were demultiplexed using the Illumina pipeline v1.9. Adapter sequences were trimmed using cutadapt v1.6 [45]. Trimmed sequences were then aligned to multiple target sequence databases, using BWA v0.7.10 (MEM algorithm with parameters ‘-v 2 –t 4’) [46]: (1) all trimmed sequences were aligned against a ribosomal RNA database to filter out residual rRNA that escaped depletion during library preparation; (2) remaining sequences were then aligned against a custom database of potential microbiome component species (see below) using BWA; (3) sequences that did not align to either the rRNA or microbiome databases were aligned to all *D. melanogaster* sequences in RepBase [47]. The remaining sequences that did not align to any of the databases above were then aligned to the *D. melanogaster* genome (BDGP5) and known transcriptome (FlyBase v5.57) using STAR v2.4.0e [48].

#### Gene expression estimation

We followed the analysis described previously [26] to compute read counts for known and novel gene models using HTSeq-count [49] with the ‘intersection-nonempty’ assignment method. Tabulated read counts for each endogenous gene present in both Baseline and ethanol-treated lines were combined and normalized across all samples using EdgeR [50]. The normalized gene expression was used in all following analyses.

#### Genetics of gene expression

For each expression feature (known and novel transcripts) we fit mixed-effect models to the normalized gene expression data corresponding to: *Y* = *S* + *W* + *T*+ *L* + *W*×*S* + *L*×*S* + *L*×*T* + *T*×*S* + *T*×*S*×*L* + *ε,* where *Y* is the observed log2 (normalized read count), *S* is sex, *W* is Wolbachia infection status, *W*×*S* is Wolbachia by sex interaction, *L* is DGRP line, *T* is treatment (ethanol-supplemented *vs* standard medium), *L*×*S* is the line by sex interaction, *L*×*T* is the line by treatment interaction, *T*×*S*×*L* is the treatment by line by sex interaction and *ε* is the residual error. We also performed reduced analyses for sexes separately (*Y* = *W* + *L* + *T*+ *L*× *T +* ε).

We identified genetically variable transcripts as those that passed a 5% FDR threshold (based on Benjamini-Hochberg corrected *P*-values [51]) for the *L, T* and *L*×*T* terms. We computed the broad sense heritabilities (*H*^2^) for each gene expression trait separately for males and females as 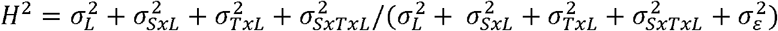, where 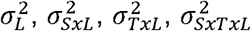 and 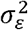 are, respectively, the among line, sex by line, treatment by line, sex by treatment by line, and within line variance components.

In addition, for all expression features that were significant for the *L*×*T* interaction term we re-analyzed data for each Treatment condition separately to identify transcripts with significant changes in expression in one or another condition, or both. We performed reduced analysis for sexes separately (*Y* = *W* + *L* + *ε*). We also calculated the Line means differences for the matching transcripts that were significant for the *L*×*T* term (*i.e.,* Line.ETOH – Line.Baseline) for males and females separately. These line means differences in expression values were used for hierarchical clustering analysis for the subset of the significant gene expression features using the JMP12 package (SAS, Cary, USA).

### Construction of correlated expression networks

The differences between the conditional means with and without ethanol for each line were used to calculate pairwise Pearson correlation coefficients for all genes that had statistically significant Benjamini-Hochberg False Discovery Rate (BH-FDR) [51] adjusted p-values (BH-FDR < 0.05) for the line-by-ethanol treatment interaction term in the linear mixed effects model (*Y = W + L + T + LxT + ε* run for each sex separately). The delta-expression correlation matrix was bi-clustered using hierarchical clustering with complete linkage agglomeration in Genesis statistical software [52]. The bi-clustered matrix was used for the center panel in each composite figure (Figs. 2 and 3). To generate the delta-expression correlation networks, the correlation matrix was filtered for associations with BH-FDR adjusted p-value < 0.05 and the top 10% of correlations based on the absolute value of the Pearson coefficient. Associations that survive the stringent filtering criteria were input into Cytoscape and clustered using the MCODE algorithm with default parameters, but with the ‘Fluff’ setting activated to capture relationships outside of auto-correlated modules [53]. The resulting modules with significantly strong intra-connectivity (cumulative MCODE score > 4) were mapped back to the correlation matrix panel based on the identity of the gene membership of each module. Genes with a statistically significant (BH-FDR < 0.05) eQTL association calculated from the expression differences were highlighted in each module within the composite figure.

### eQTL mapping

eQTLs were mapped to differences in expression between baseline and ethanol treatment of genes with a statistically significant line-by-treatment (*L*x*T*) term from the linear mixed effects model run for each sex separately as previous described [25,26]. Briefly, normalized FPKM values were adjusted for *Wolbachia* infection status, chromosomal inversions, population structures organized based on top 10 principal components using Best Linear Unbiased Predictor (BLUP) using the R package *lmerTest.* Covariate adjusted expression differences were used as phenotype for eQTL mapping using PLINK (v1.90). Association *P-*values generated by the PLINK *t*-tests were compared to the empirical FDR threshold calculated by dividing the number of expected associations under the null hypothesis generated from 100 permutations at a false discovery rate of 0.05 by the observed number of associations at the same threshold to determine statistical significance. Associations were further filtered for independence using forward model selection, as previously described [25,26], by iteratively adding single eQTLs, starting with the smallest *P*-values, to an additive association model such that the conditional *P-*value for the last added eQTL is no more than 1E-5.

### Association networks

To build association networks using the variants identified from eQTL mapping, pairwise eQTL associations between genes that either contain the variant or are within 1000 bp up- or down-stream of the variant and genes with statistically significant line-by-treatment (*LxT*) were added to the most recent version (fb_2021_05) of the database of known genetic and protein-protein or RNA-protein physical interactions from the FlyBase repository and visualized in Cytoscape. The resulting network of associations were filtered to contain (i) genes that are part of the eQTL associations, and (ii) genes that have at least 5 genetic or physical interactions to genes that are part of the eQTL associations within one interaction distance (one edge). The filtered interaction network was modularized using MCODE algorithm with default settings but with ‘fluff’ activated [53]. Genes that are part of the individual modules were used for Gene Ontology Enrichment analysis. Statistically significant (FDR < 0.05 in the Overrepresentation Test), highly specialized terms containing the largest number of genes from the input from each module were used for functionally labeling each module. For the glutathione metabolism inlet in the female interaction network in Fig. 3, *GstE* family of genes were added to the *GstD* tri-gene cluster based on semantic similarity.

### Analysis of activity and sleep phenotypes

Flies were reared on standard medium and medium supplemented with 10% (v/v) ethanol and placed in standard food collection vials overnight. The next day, mated females and males from both treatment conditions were placed in Drosophila Activity Monitor (DAM) tubes (TriKinetics, Waltham, MA) that contained agar supplemented with 5% sucrose at one end, and a small piece of yarn at the other end. Flies were placed in a 25°C incubator on a 12-hour light-dark cycle and their activity and sleep data were recorded using the DAMSystem (Trikinetics). Raw data from the DAMSystem (TriKinetics) were uploaded to ShinyR-DAM [54] and resulting output data were parsed by sleep/activity phenotype for analysis. Sleep was defined as at least 5 minutes of inactivity and only data from flies that survived the entire testing period (2-9 days of the fly lifespan) were retained for analysis. Sleep and activity data were analyzed using the PROC MIXED command (Type III Analysis of Variance (ANOVA)) within SAS version 9.04 (Cary, NC) according to the model *Y* = *μ* + *T* + *L* + *S* + *TxL* + *TxS* + *LxS* + *TxLxS* + *Rep(TxLxS)* + *ε*, where *T* is the fixed effect of treatment (ethanol medium, standard medium), *L* is the fixed effect of line (RAL_177, RAL_208, RAL_367, RAL_555, RAL_705, RAL_730), *S* is the fixed effect of Sex (male, female), *Rep(TxLxS)* is the random effect of replicate, and *ε* is the residual variance. Reduced Type III ANOVAs (*Y* = *μ* + *T* + *L* + *TxL* + *Rep(TxL)* + *ε)* were also performed by Sex.

## Availability of data and materials

RNA sequencing data have been deposited in the GEO database under accession number GSE186240. Raw DAM data are on the github repository: https://github.com/rebeccamacpherson/DAM_raw_data_DEV_ETOH_DGRP

## Authors’ contributions

TVM, TFCM and RRHA designed research; TVM performed the transcriptional profiling studies, RAM performed the behavioral studies; TVM, VS and RAM analyzed data; TVM, RRHA and TFCM wrote the paper.

## Acknowledgments

We thank Gunjan Arya, Lavanya Turlapati and Genevieve St. Armour for technical assistance. This work was supported by grants from the National Institutes of Health (AA016560 and DA041613) to RRHA and TFCM.

## Supplementary material

**Figure S1.** Correlations between variation in expression of snoRNAs after chronic exposure to ethanol and variation in expression of their host genes. *CycE* is included since it is a hub gene in a genetic network associated with variation in ethanol-induced variation in development time and viability [16], and *Myc* is included as it is associated with ribosome biogenesis and has been implicated as a regulator of *Uhg4* [38]. The graphs on the right illustrate examples of scatter plots of the correlations between expression of *Uhg4* and several snoRNAs.

**Figure S2.** Correlations between ethanol-induced variation in expression of *snoRNA* host genes and variation in expression of *CycE.* The scatter plots illustrate examples of correlations between expression of *CycE* and several snoRNA host genes.

**Additional File 1.** DGRP lines used for transcriptional profiling.

**Additional File 2.** Three-way mixed model ANOVA for differences in transcript abundances between flies reared on regular and ethanol supplemented medium for sexes combined. The main effects are treatment (fixed, reared on standard vs. ethanol supplemented medium), Sex (fixed, males and females) and DGRP Line (random). All interactions with Line are random, the others are fixed.

**Additional File 3.** ANOVA for differences in transcript abundances between flies reared on regular and ethanol supplemented medium for (A) females and (B) males.

**Additional File 4.** Genes with significant *LxT* interactions in transcript abundances. (A) Results of female-only mixed-effect models for all expressed gene profiles, including alignment bias estimates. (B) Results of male-only mixed-effect models for all expressed gene profiles, including alignment bias estimates. (C) Genes with significant *LxT* interactions common to females and males and that are female-specific or male-specific. (D) Enrichment analyses for genes with significant *LxT* interactions in transcript abundances in females. (E) Enrichment analyses for genes with significant *LxT* interactions in transcript abundances in males. (F) Enrichment analyses for genes with significant *LxT* interactions in transcript abundances in both sexes. (G) Enrichment analyses for genes with significant *LxT* interactions in transcript abundances in females only. (H) Enrichment analyses for genes with significant *LxT* interactions in transcript abundances in males only. (I) Human orthologs for genes with significant *LxT* interactions in transcript abundances.

**Additional File 5. (**A) eQTL mapping and model selection results for genes with a statistically significant *L*x*T* ANOVA term in females and males. (B) Genes containing eQTLs (in the gene body or within a 1,000 bp window of the eQTL) in females and males. Intergenic: > 1,000 bp from any gene body.

**Additional File 6.** snoRNAs with altered gene expression following chronic exposure to ethanol during development and their host genes. snoRNAs that occur in clusters without intervening genes are in bold font and square brackets.

**Additional File 7.** Mixed model ANOVAs for sleep and activity phenotypes. Line, Sex, and Treatment are fixed effects, Replicate (Rep) is random. “df”: degrees of freedom; “SS”: Type III Sums of Squares; “MS”: Mean square.

